# Recalibrating the Epigenetic Clock: Implications for Assessing Biological Age in the Human Cortex

**DOI:** 10.1101/2020.04.27.063719

**Authors:** Gemma L Shireby, Jonathan P Davies, Paul T Francis, Joe Burrage, Emma M Walker, Grant W A Neilson, Aisha Dahir, Alan J Thomas, Seth Love, Rebecca G Smith, Katie Lunnon, Meena Kumari, Leonard C Schalkwyk, Kevin Morgan, Keeley Brookes, Eilis J Hannon, Jonathan Mill

**Affiliations:** University of Exeter Medical School, University of Exeter, Exeter, UK; Wolfson Centre for Age-Related Diseases, King’s College London, London, UK; Institute of Neuroscience, Newcastle University, Newcastle Upon Tyne, UK; Dementia Research Group, Institute of Clinical Neurosciences, School of Clinical Sciences, University of Bristol, Bristol, UK; Institute for Social and Economic Research, University of Essex, Colchester, UK; School of Life Sciences, University of Essex, Colchester, UK; Human Genetics Group, University of Nottingham, Nottingham, UK; School of Science & Technology, Nottingham Trent University, Nottingham, UK

**Author notes:** These authors contributed equally (Eilis J Hannon, Jonathan Mill).

**Keywords:** Cortex, age, ageing, disease, epigenetic clock, DNA methylation, post-mortem

## Abstract

Human DNA-methylation data have been used to develop biomarkers of ageing - referred to ‘epigenetic clocks’ - that have been widely used to identify differences between chronological age and biological age in health and disease including neurodegeneration, dementia and other brain phenotypes. Existing DNA methylation clocks are highly accurate in blood but are less precise when used in older samples or on brain tissue. We aimed to develop a novel epigenetic clock that performs optimally in human cortex tissue and has the potential to identify phenotypes associated with biological ageing in the brain. We generated an extensive dataset of human cortex DNA methylation data spanning the life-course (n = 1,397, ages = 1 to 104 years). This dataset was split into ‘training’ and ‘testing’ samples (training: n = 1,047; testing: n = 350). DNA methylation age estimators were derived using a transformed version of chronological age on DNA methylation at specific sites using elastic net regression, a supervised machine learning method. The cortical clock was subsequently validated in a novel human cortex dataset (n = 1,221, ages = 41 to 104 years) and tested for specificity in a large whole blood dataset (n = 1,175, ages = 28 to 98 years). We identified a set of 347 DNA methylation sites that, in combination optimally predict age in the human cortex. The sum of DNA methylation levels at these sites weighted by their regression coefficients provide the cortical DNA methylation clock age estimate. The novel clock dramatically out-performed previously reported clocks in additional cortical datasets. Our findings suggest that previous associations between predicted DNA methylation age and neurodegenerative phenotypes might represent false positives resulting from clocks not robustly calibrated to the tissue being tested and for phenotypes that become manifest in older ages. The age distribution and tissue type of samples included in training datasets need to be considered when building and applying epigenetic clock algorithms to human epidemiological or disease cohorts.

## Introduction

Advancing age is associated with declining physical and cognitive function, and is a major risk factor for many human brain disorders including dementia and neurodegenerative disease (Harper, 2014; Sierra, 2019). Understanding the biological mechanisms involved in ageing will be a critical step towards preventing, slowing or reversing age-associated phenotypes. Due to the substantial inter-individual variation in age-associated phenotypes, there is considerable interest in identifying robust biomarkers of ‘biological’ age, a quantitative phenotype that is thought to better capture an individuals’ risk of age-related outcomes than actual chronological age (Jylhävä *et al*., 2019). Several data modalities have been used to generate estimates of biological age; these include measures of physical fitness (e.g. muscle strength) (Sosnoff and Newell, 2006), cellular phenotypes (e.g. cellular senescence) (Baker *et al*., 2011) and genomic changes (e.g. telomere length) (Jylhävä *et al*., 2017; Sanders and Newman, 2013).

Epigenetic mechanisms act to regulate gene expression developmentally via chemical modifications to DNA and histone proteins (Bernstein *et al*., 2007), conferring cell-type-specific patterns of gene expression and differing markedly between tissues and celltypes (Mendizabal and Yi, 2016). There has been recent interest in the dynamic changes in epigenetic processes over the life course, and a number of ‘epigenetic clocks’ based on one specific epigenetic modification - DNA methylation (DNAm) - have been developed that appear to be highly predictive of chronological age (Campisi and Vijg, 2009; Horvath, 2013; Horvath *et al*., 2012, 2018; Knight *et al*., 2016; Oberdoerffer and Sinclair, 2007; Simpkin *et al*., 2017). The landmark DNAm clock was developed by Horvath (Horvath, 2013), who applied elastic net regression to Illumina DNAm array data from a large number of samples derived from a range of tissues (n = ~ 8,000 across 51 tissue and cell types), and generated a predictor based on DNAm at 353 CpG sites that is highly predictive of chronological age (Horvath, 2013). Given that changes in DNAm are known to index exposure to certain environmental risk factors for diseases of old age (for example, tobacco smoking) (Elliott *et al*., 2014; Sugden *et al*., 2019), and variable DNAm is robustly associated with a number of age-associated disorders (Chouliaras *et al*., 2018; Chuang *et al*., 2017; Smith *et al*., 2016), there has been interest in the hypothesis that DNAm clocks might robustly quantify variation in biological age. Horvath’s DNAm age clock, for example, has been widely applied to identify accelerated epigenetic ageing - where DNAm age predictions deviate from chronological age such that individuals appear older than they really are - in the context of numerous health and disease outcomes (Horvath and Ritz, 2015; Levine *et al*., 2015; Marioni *et al*., 2015; McCartney *et al*., 2018). Since age is a major risk factor for dementia and other neurodegenerative brain disorders, there is particular interest in the application of epigenetic clock algorithms to these phenotypes, especially as differential DNAm has been robustly associated with diseases including Alzheimer’s disease and Parkinson’s disease (Lunnon *et al*., 2014; Smith *et al*., 2016; Yu *et al*., 2015). Recent studies have reported an association between accelerated DNAm age and specific markers of Alzheimer’s disease neuropathology in the cortex (e.g. neuritic plaques, diffuse plaques and amyloid-β load) (Levine *et al*., 2015, 2018). Furthermore, among individuals with Alzheimer’s disease, DNAm age acceleration is associated with declining global cognitive functioning and deficits in episodic and working memory (Levine *et al*., 2015, 2018).

A major strength of existing epigenetic clocks is that they work relatively well across different types of sample; the Horvath multi-tissue clock, for example, can accurately predict age in multiple tissues across the life-course. However, as with any predictor, the composition of the training data used to develop the clock influences the generality of the model. To date, there has been limited research comparing the prediction accuracy and potential bias of existing clock algorithms across different tissues and ages. Recent analyses have highlighted potential biases when using Horvath’s clock in older samples (>~60 years) and in samples derived from certain tissues, especially the central nervous system (El Khoury *et al*., 2019). This is important for the interpretation of studies of possible relationships between accelerated epigenetic age and age-related diseases affecting the human brain (e.g. dementia and neurodegenerative phenotypes); reported associations between accelerated DNAm age and disease may actually be a consequence of fitting a suboptimal predictor to available datasets. Potential confounders include differential changes in DNAm with age across tissues, and the age distribution of the samples used to train existing classifiers. Resolution of these biases requires the construction of specific DNAm clocks developed using data generated on the relevant tissue-type and including broad representation of the age spectrum they will be used to interrogate. Recently, a number of tissue-specific DNA methylation clocks have been described, including clocks designed for whole blood (Hannum *et al*., 2013; Zhang *et al*., 2019), muscle (Voisin *et al*., 2019), bone (Gopalan *et al*., 2019) and paediatric buccal cells (McEwen *et al*., 2019). Importantly, although these DNAm age estimators have increased predictive accuracy within the specific tissues in which they were built, they lose this precision when applied to other tissues (El Khoury *et al*., 2019).

We describe the development of a novel DNAm clock that is specifically designed for application in DNA samples isolated from the human cortex and is accurate across the lifespan including in tissue from elderly donors. We demonstrate that our clock outperforms existing DNAm-based predictors developed for other tissues, minimising the potential for spurious associations with ageing phenotypes relevant to the brain.

## Materials and methods

### Datasets used to develop the novel cortical DNAm age clock

To develop and characterise our cortical DNAm age clock (“DNAmClock_Cortical_”) we collated an extensive collection of DNAm data from human cortex samples (see **Supplementary Table 1**), complementing datasets generated by our group (http://www.epigenomicslab.com) with publicly available datasets downloaded from the Gene Expression Omnibus (GEO; https://www.ncbi.nlm.nih.gov/geo/) (Jaffe *et al*., 2016; De Jager *et al*., 2014; Lunnon *et al*., 2014; Pidsley *et al*., 2014; Smith *et al*., 2018, 2019; Wong *et al*., 2019) (see **Supplementary Table 1**). In each of these datasets DNAm was quantified across the genome using the Illumina 450K DNA methylation array which covers >450,000 DNA methylation sites as previously described (Pidsley *et al*., 2013). To optimise the performance of the DNAmClock_Cortical_ and to avoid reporting over-fitted statistics, the samples were split into a “training” dataset (used to determine the DNAm sites included in the model and their weighted coefficients) and a “testing” dataset (used to profile the performance of the proposed model). To reduce the effects of experimental batch in our model, we maximised the number of different datasets included in the training data by combining the ten cohorts and randomly assigning individuals within them to either the training or testing dataset in a 3:1 ratio (**Table 1**). In total, our training dataset (age range = 1-108 years, median = 57 years; see **Supplementary Fig. S1**) comprised DNAm data from 1,047 cortex samples (derived from 832 donors) and our testing dataset (age range = 1-108 years, median = 56 years; see **Supplementary Fig. S1**) comprised DNAm data from 350 cortex samples (derived from 323 donors). Individuals with a diagnosis of Alzheimer’s disease and other major neurological phenotypes were excluded from our analysis given the previous associations between them and deviations in DNAm age (Levine *et al*., 2015, 2018).

**Table 1:** Sample characteristics of the training (cortex), testing (cortex), validation (cortex) and whole blood datasets used in the development and evaluation of our novel cortical DNA methylation clock.

| Dataset | Age (years) |  |  |  | Sex (number) |  | Illumina<br>BeadCHIPArray | References for data |
| --- | --- | --- | --- | --- | --- | --- | --- | --- |
|  | N | Mean | Median | Range | Female | Male |  |  |
| Training | 1047 | 56.53 | 57 | 1-108 | 362 | 685 | 450K | (Jaffe *et al.*, 2016; De Jager *et al.*, 2014; Lunnon *et al.*, 2014; Pidsley *et al.*, 2014; Smith *et al.*, 2018, 2019; Wong *et al.*, 2019) |
| Testing | 350 | 55.87 | 56 | 1-108 | 144 | 206 | 450K | (Jaffe *et al.*, 2016; De Jager *et al.*, 2014; Lunnon *et al.*, 2014; Pidsley *et al.*, 2014; Smith *et al.*, 2018, 2019; Wong *et al.*, 2019) |
| Validation | 1221 | 83.49 | 84 | 41-104 | 577 | 644 | EPIC | - |
| Blood | 1175 | 57.96 | 59 | 28-98 | 686 | 489 | EPIC | Hannon et al. (2018) |

### Cortex validation dataset

An independent “validation” cortical dataset was generated using post-mortem occipital (OCC) and prefrontal cortex (PFC) samples from the Brains for Dementia Research (BDR) cohort. BDR was established in 2008 and is a UK-based longitudinal cohort study with a focus on dementia research (Francis *et al*., 2018) coordinated by a network of six dementia research centres based around the UK. Post-mortem brains underwent full neuropathological dissection, sampling and characterisation using a standardised protocol (Bell *et al*., 2008; Samarasekera *et al*., 2013). DNA was isolated from cortical tissue samples using the Qiagen AllPrep DNA/RNA 96 Kit (Qiagen, cat no.80311) following tissue disruption using BeadBug 1.5 mm Zirconium beads (Sigma Aldrich, cat no.Z763799) in a 96-well Deep Well Plate (Fisher Scientific, cat no.12194162) shaking at 2500rmp for 5 minutes. Genome-wide DNA methylation was profiled using the Illumina EPIC DNA methylation array (Illumina Inc), which interrogates >850,000 DNA methylation sites across the genome (Moran *et al*., 2016). After stringent data quality control (see **below**) the final validation dataset consisted of DNAm estimates for 800,916 DNAm sites profiled in 1,221 samples (632 donors; 610 PFC; 611 OCC; see **Table 1** for more details). This dataset consists of predominantly elderly samples (age range = 41-104 years, median = 84 years; see **Supplementary Fig. S1**).

### Whole blood dataset

We recently generated DNAm data from whole blood obtained from 1,175 individuals (age range = 28-98 years; median age = 59 years; see **Table 1** for more details) included in the UK Household Longitudinal Study (UKHLS) (https://www.understandingsociety.ac.uk/) (Hannon *et al*., 2018). The UKHLS was established in 2009 and is a longitudinal panel survey of 40,000 UK households from England, Scotland, Wales and Northern Ireland (Buck and McFall, 2011). For each participant, non-fasting blood samples were collected through venipuncture; these were subsequently centrifuged to separate plasma and serum, and samples were aliquoted and frozen at −80°C. DNAm data were generated using the Illumina EPIC DNA methylation array as described previously ((Hannon *et al*., 2018). After stringent QC (see **below**) the whole blood dataset consisted of data for 857,071 DNAm sites profiled in 1,175 samples (Hannon *et al*., 2018).

### DNA methylation data pre-processing

Unless otherwise reported, all statistical analysis was conducted in the R statistical environment (version 3.5.2; https://www.r-project.org/). Raw data for all datasets were used, prior to any QC or normalisation, and processed using either the *wateRmelon* (Pidsley *et al*., 2013) or *bigmelon* (Gorrie-Stone *et al*., 2019) packages. Our stringent QC pipeline included the following steps: (1) checking methylated and unmethylated signal intensities and excluding poorly performing samples; (2) assessing the chemistry of the experiment by calculating a bisulphite conversion statistic for each sample, excluding samples with a conversion rate <80%; (3) identifying the fully methylated control sample was in the correct location (where applicable); (4) multidimensional scaling of sites on the X and Y chromosomes separately to confirm reported sex; (5) using the 65 SNP probes present on the Illumina 450K array and 59 on the Illumina EPIC array to confirm that matched samples from the same individual (but different brain regions) were genetically identical and to check for sample duplications and mismatches; (6) use of the *pfilter()* function in *wateRmelon* to exclude samples with >1 % of probes with a detection P value > 0.05 and probes with >1 % of samples with detection P value > 0.05; (8) using principal component analysis on data from each tissue to exclude outliers based on any of the first three principal components; (9) removal of cross-hybridising and SNP probes (Chen *et al*., 2013). The subsequent normalisation of the DNA methylation data was performed using the *dasen()* function in either *wateRmelon* or *bigmelon* (Gorrie-Stone *et al*., 2019; Pidsley *et al*., 2013).

### Deriving a novel cortical DNAm age classifier

To build the DNAmClock_Cortical_ we implemented an elastic net (EN) regression model, using the methodology described by Horvath (2013). The EN model is designed for high dimensional datasets with more features than samples and where the features are potentially highly correlated (Zou and Hastie, 2005). As part of the methodology, the model selects the subset of features (i.e. DNAm sites) that cumulatively produce the best predictor of a provided outcome. EN was implemented in the R package *GLMnet* (Friedman *et al*., 2010). It uses a combination of Ridge and LASSO (Least Absolute Shrinkage and Selection Operator) regression. Ridge regression penalises the sum of squared coefficients and has an (alpha) parameter of zero. LASSO regression penalises the sum of the absolute values of the coefficients and has an ***α*** parameter of one. EN is a convex combination of ridge and LASSO and, therefore, the elastic net ***α*** parameter was set to 0.5. The lambda value (the shrinkage parameter) was derived using 10-fold cross-validation on the training dataset (lambda = 0.0178). DNAm probes included in the analysis were limited to sites which were present on both the Illumina EPIC and Illumina 450K arrays, with no missing values across the training datasets (n probes = 383547). Previous analyses have shown that the relationship between DNAm age (predicted age from epigenetic age estimators) and chronological age is logarithmic between 0-20 years and linear from 20 years plus (Horvath, 2013). Our data revealed a similar pattern and therefore chronological age was transformed (**Supplementary Fig. S2**). A transformed version of chronological age was regressed on DNAm levels at all included DNAm sites.

### Implementing DNAm Age prediction

We applied the DNAmClock_Cortical_ (comprising 347 DNAm sites) to the testing, validation and whole blood DNAm datasets. We then compared its performance to a number of existing DNAm clocks: Horvath’s original multi-tissue clock (“DNAmClock_Multi_”; 353 DNAm sites) (Horvath, 2013), Zhang’s EN blood-based DNAm clock (“DNAmClock_Blood_”: 514 DNAm sites) (Zhang *et al*., 2019) and Levine’s ‘pheno age’ DNAm Clock (“DNAmClock_Pheno_”; 513 DNAm sites) (Levine *et al*., 2018). Briefly, to predict DNAm age using the DNAmClock_Multi_ we applied the *agep()* function in *wateRmelon* (Pidsley *et al*., 2013). Although this function does not contain the custom normalisation method applied at the DNAm age calculator website (https://DNAmClock.genetics.ucla.edu/), both methods work similarly in brain and blood studies, providing the data have been pre-processed adequately (El Khoury *et al*., 2019). To predict age using the DNAmClock_Pheno_ (Levine *et al*., 2018), we also applied the *agep()* function, inputting a vector of the coefficients and the intercept using the data provided in the supplementary material of Levine et al’s manuscript. To predict DNAm age with the DNAmClockblood, we used the authors’ published code (available on GitHub https://github.com/qzhang314/DNAm-based-age-predictor) (Zhang *et al*., 2019).

### Determining the predictive accuracy of different DNAm clocks

DNAm age was estimated in the testing dataset (n = 350), validation dataset (n = 1221) and whole blood dataset (n = 1175) using the DNAmClock_Cortical_, DNAmClock_Multi_, DNAmClock_Blood_ and the DNAmClock_Pheno_. To compare and evaluate the predictive accuracy of these DNAm age predictors, estimates were assessed using two measures: Pearson’s correlation coefficient (r; a measure indicating the strength of the linear relationship between the actual (chronological age) and predicted (DNAm age) variables) and the root mean squared error (RMSE; square root of the mean differences between the actual and predicted variables) which quantifies the precision of the estimator.

### Analysis against age

To test associations between DNAm age and chronological age, we fitted regression models to each dataset. As a subset of donors included in the testing and validation datasets had data from multiple cortical regions, we used mixed effects linear regression models, implemented with the *lme4* and *lmerTest* packages, where DNAm age was regressed against chronological age as a fixed effect and individual was included as a random effect. In the blood cohort, as there was only one sample per individual, we applied standard linear regression models. A second regression model was also fitted which additionally tested for associations with an age-squared term.

### Data availability

The datasets used for the training and testing samples are available for download from GEO (https://www.ncbi.nlm.nih.gov/geo/) using the following accession numbers: GSE74193; GSE59685; GSE80970; GPL13534 and GSE43414. The validation data are available from the authors upon reasonable request. The whole blood DNA methylation data are available upon application through the European Genome-Phenome Archive under accession code EGAS00001001232. Analysis scripts used in this manuscript are available on GitHub (https://github.com/qzhang314/DNAm-based-age-predictor).

## Results

### Existing DNAm clock algorithms work sub-optimally in the human cortex, systematically underestimating age in elderly individuals

The performance of DNAm clocks is influenced by the characteristics (e.g. specific tissue type and age range) of the training data used to build the prediction algorithm. Applying predictors to datasets that differ in terms of these characteristics may lead to biases when estimating DNAm age, and confound phenotypic analyses using these variables (El Khoury *et al*., 2019). We found that existing DNAm clocks (i.e. the DNAmClock_Multi_ (Horvath, 2013) the DNAmClock_Blood_ (Zhang *et al*., 2019) and the DNAmClock_Pheno_ (Levine *et al*., 2018)) do not perform optimally in human cortex tissue (**Supplementary Fig. S3**), with notable differences between derived DNAm age and actual chronological age (i.e. the derived values do not lie along the y = x line, see **Fig. 1**). In our testing dataset (n = 350 cortex samples; age range = 1 - 108 years; median age = 57 years), the DNAmClock_Multi_ systematically overestimated DNAm age in individuals over ~60, and systematically underestimated it in individuals below ~60 years (**Fig. 1A**(**ii**) and **Fig. 2A**(**ii**)). In the elderly group, around 80% of samples had lower predicted DNAm ages than their actual chronological age. These deviations were also observed when looking at the mean differences between actual age and predicted DNAm age (referred to as Δ (delta) age), such that Δ age was positive for younger ages and vice versa for the elderly group (**Supplementary Fig. S4A**). Use of the DNAmClock_Blood_ produced even more pronounced systematic underestimation of DNAm age in adults, starting around 30 years (**Fig. 1A**(**iii**) and **Fig. 2A**(**iii**)), and this trend was mirrored for Δ age (see **Supplementary Fig. S4A**). Finally, the DNAmClock_Pheno_ severely under predicted age in the cortex, with 100% of samples being assigned a lower DNAm age than the actual chronological age (**Fig. 1A**(**iv**)**, Fig. 2A**(**iv**) **and Supplementary Fig. S4A**(**iv**)). Similar biases in age prediction were seen in our validation cohort (n = 1,221 cortex samples; age range = 41 years to 104 years; mean age = 83.49 years), confirming the systematic underestimation of DNAm age in elderly donors (see **Fig. 1B** and **Fig. 2B**). As with the other clocks, Δ age captured these biases, with particularly poor performance evident when applying the DNAmClock_Pheno_ and the DNAmClock_Blood_ to this dataset, in which Δ age was consistently below zero (where zero would represent perfect prediction; see **Supplementary Fig. S4B**).

**Figure 1:**
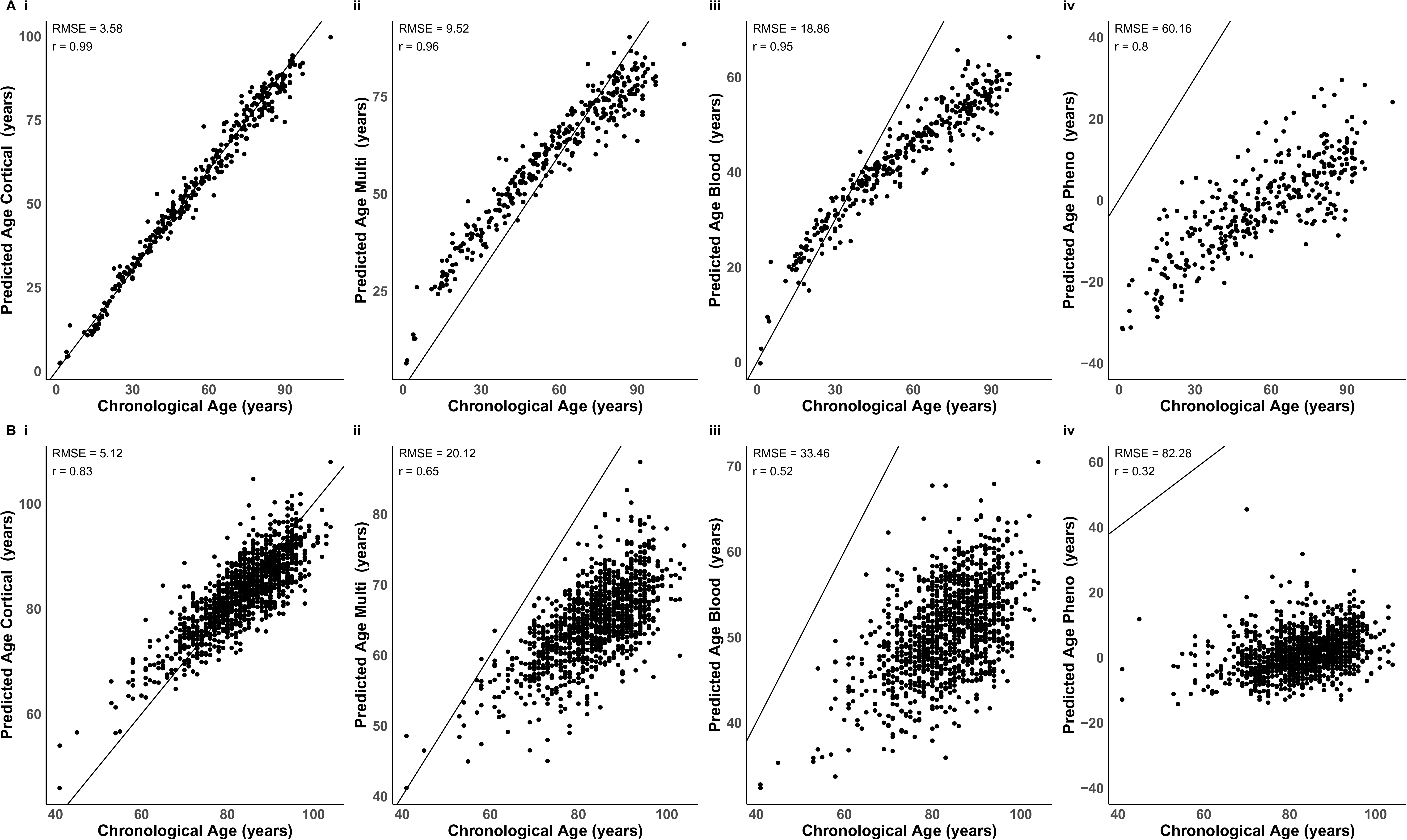
Comparison of chronological age with DNA methylation age derived using four DNA methylation age clocks. Shown are comparisons of chronological age with predicted age in (**A**) the testing dataset (n = 350 cortical samples) and (**B**) the validation dataset (n = 1221 cortical samples). DNAm age was predicted using four DNA methylation age clocks: (**i**) our novel DNAmClock_Cortical_; (**ii**) Horvath’s DNAmClock_Multi_; (**iii**) Zhang’s DNAmClock_Blood_ and (**iv**) Levine’s DNAmClock_Pheno_. The x-axis represents chronological age (years) and the y-axis represents predicted age (years). Each point on the plot represents an individual sample. Our cortical clock out-performed the three alternative DNAm clocks across all accuracy statistics. DNA methylation age estimates derived using the DNAmClock_Multi_ (**A**(**ii**) testing and **B**(**ii**) validation) and the DNAmClock_Blood_ (**A**(**iii**) testing and **B**(**iii**) validation) appear to have a non-linear relationship with chronological age. *DNAmClock_Cortical_ = Cortical DNA methylation age Clock; DNAmClock_Multi_ = Multi-tissue DNA methylation age clock; DNAmClock_Blood_ = Blood DNA methylation age clock and DNAmClock_Pheno_ = Pheno Age DNA methylation age clock.

**Figure 2:**
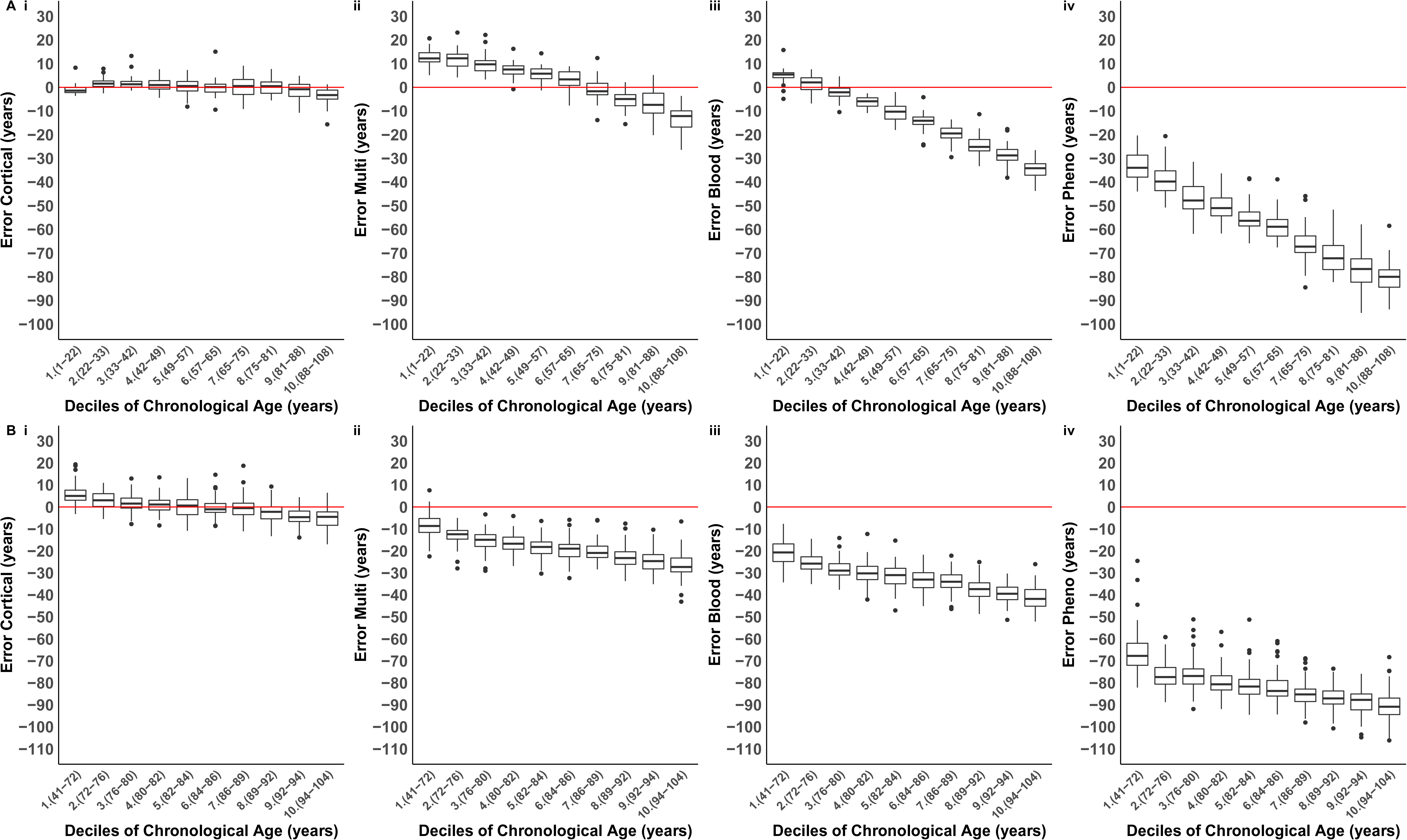
The cortical DNA methylation age clock has elevated accuracy in human cortex samples across the lifespan. Shown is the distribution of the error (DNA methylation age - chronological age) for each age decile in (**A**) the testing dataset (n = 350 cortical samples) and (**B**) the validation dataset (n = 1221 cortical samples) for each of the four DNA methylation age clocks: (**i**) our novel DNAmClock_Cortical_; (**ii**) Horvath’s DNAmClock_Multi_; (**iii**) Zhang’s DNAmClock_Blood_ and (**iv**) Levine’s DNAmClock_Pheno_. Deciles were calculated by assigning chronological age into ten bins and are represented along the x-axis by the numbers one to ten, followed by brackets which display the age range included in each decile. The ends of the boxes are the upper and lower quartiles of the errors, the horizontal line inside the box represents the median deviation and the two lines outside the boxes extend to the highest and lowest observations. Outliers are represented by points beyond these lines. The red horizontal line represents perfect prediction (zero error). Our novel DNAmClock_Cortical_(**A**(**i**) testing and **B**(**i**) validation) consistently had the smallest error across the age groups, shown by the tightness of the boxplot distributions along the zero-error line. The DNAmClock_Multi_ over-predicted younger ages (deciles 1-5 in **A**(**ii**)), shown by boxplots distributions which are above the zero-error line, and under predicted older ages (deciles 8-10 in **A**(**ii**) and deciles 1-10 in **B**(**ii**)), shown by boxplot distributions below the zero-error line. The DNAmClock_Blood_ (**A**(**iii**) testing and **B**(**iii**) validation) and the DNAmClock_Pheno_ (**A**(**iv**) testing and **B**(**iv**) validation) consistently underpredicted age, with underprediction of DNA methylation age increasing with chronological age. *DNAmClock_Cortical_ = Cortical DNA methylation age Clock; DNAmClock_Multi_ = Multi-tissue DNA methylation age clock; DNAmClock_Blood_ = Blood DNA methylation age clock and DNAmClock_Pheno_ = Pheno Age DNA methylation age clock

### Developing a novel DNAm clock for the human cortex based on 347 DNA methylation sites

The composition of the training data used to build a predictor can influence the generality of the model. Therefore, to alleviate the biases observed when applying existing DNAm clocks to data generated on elderly human cortex samples, we focussed on building a DNAm clock using relevant tissue samples from donors that spanned a broad range of ages and included a large number of samples from elderly donors (**Supplementary Fig. S1**). We developed our novel cortical DNAm clock (DNAmClock_Cortical_) using an elastic net regression, regressing chronological age against DNAm levels across 383,547 sites quantified 1,047 cortex samples (see **Methods**). This approach identified a set of 347 DNAm sites which in combination optimally predict age in the human cortex. The sum of DNAm levels at these sites weighted by their regression coefficients provides the DNAmClock_Cortical_ age estimate (see **Supplementary Table 2**). Of note, the majority of sites selected for our cortex clock were novel and not present in existing DNAm clock algorithms; only 5 of the sites overlap with the DNAmClock_Multi_ (composed of 353 DNAm sites), 15 with the DNAmClock_Blood_ (comprising 514 DNAm sites), and 5 with the DNAmClock_Pheno_ (comprising 513 DNAm sites) (see **Supplementary Table 3**).

### Increased prediction accuracy of the novel cortex clock in cortical tissue compared to existing DNAm clocks

We used the DNAmClock_Cortical_ to estimate DNAm age in both the testing (n = 350 cortex samples) and validation (n = 1221 cortex samples) datasets, and comparing the estimates to those derived using DNAmClock_Multi_, DNAmClock_Blood_ and DNAmClock_Pheno_. The DNAmClock_Cortical_ predicted age accurately in the testing dataset and there was a strong correlation between DNAm age and age (r = 0.99; **Table 2** and **Fig. 1A**(**i**)). In the validation dataset, which consisted predominantly of elderly samples, our clock also performed well and was highly correlated with age (r = 0.83), outperforming DNAmClock_Multi_ (r = 0.65), DNAmClock_Blood_ (r = 0.52), and DNAmClock_Pheno_ (r = 0.32) (see **Table 2**; **Fig. 1B**(**i**)). The most striking differences were in the accuracy of the DNAmClock_Cortical_ in comparison to previously developed DNAm clocks; it outperformed the three other clocks we tested across all accuracy statistics in both cortical datasets (**Table 2**). The biggest differences in accuracy can be seen in the validation dataset (see **Fig. 1B**; **Fig. 2B** and **Supplementary Fig. S4B**), in which the RMSE was 15 years more accurate when using the DNAmClock_Cortical_ (RMSE: 5 years) than the DNAmClock_Multi_ (RMSE: 20 years), 28 years more accurate than the DNAmClock_Blood_ (RMSE: 33 years) and 77 years more accurate than the DNAmClock_Pheno_ (RMSE: 82 years). The DNAmClock_Pheno_ was consistently the most inaccurate at estimating age in the cortical data sets (RMSE: testing 60 years; validation 82 years), followed by DNAmClock_Blood_ (RMSE: testing 19 years; validation 33 years) and the DNAmClock_Multi_ (RMSE: testing: 10 years; validation 20 years) (see **Table 2** for more details).

**Table 2 –.** Our novel cortex clock outperforms existing DNA methylation age algorithms in human cortex samples. Accuracy statistics between DNAm age estimates and chronological age using our novel cortical clock, Horvath’s multitissue clock (Horvath, 2013), Zhang’s elastic net blood clock (Zhang et al., 2019) and Levine’s Pheno Age clock (Levine et al., 2018) in both the testing (n = 350 cortical samples) and validation (n = 1221 cortical samples) datasets. RMSE = root mean squared error (years). MAD = mean absolute deviation (years).

| Testing dataset (n =350) |  |  |  |  | Validation dataset (n = 1221) |  |  |  |
| --- | --- | --- | --- | --- | --- | --- | --- | --- |
|  | Cortical Clock | Multi-tissue Clock | Blood Clock | Pheno Age Clock | Cortical Clock | Multi-tissue Clock | Blood Clock | Pheno Age Clock |
| **Correlation (r)** | 0.99 | 0.96 | 0.95 | 0.8 | 0.83 | 0.65 | 0.52 | 0.32 |
| **RMSE (years)** | 3.58 | 9.52 | 18.86 | 60.16 | 5.12 | 20.12 | 33.46 | 82.28 |

### The relationship between age and DNAmClock plateaus in old age

By definition, DNAm age is correlated with chronological age, meaning age is a potential confounder for analyses of Δ age; not adequately controlling for age increases the likelihood that false positive associations will be identified (El Khoury *et al*., 2019). To assess associations between DNAm age and chronological age we used a mixed effects regression model (see **Methods**) and found that estimates from all four DNAm age clocks were significantly associated with age in the testing dataset (Bonferroni P < 0.008, see **Table 3**). Many studies of Δ m age in health and disease control for age by using a linear model to regress out its effect (Marioni *et al*., 2015; McKinney *et al*., 2018) although one of the assumptions of this approach is that the prediction accuracy of the DNAm clock is consistent across the life course. If the accuracy varies non-linearly with chronological age, then simply including age as a linear covariate in association analyses will not sufficiently negate the confounding effect of age. We therefore sought to formally test the extent to which the prediction accuracy of the four clocks correlates with age by including an age squared term in the regression model. In the testing dataset all four clocks had a significant age squared term (**Table 3**), indicating that their predictive accuracy varies as a function of age. Specifically, all clocks were associated with a plateau where the difference between DNAm age and chronological age becomes larger as actual age increases (**Fig. 2**). Importantly, however, the coefficient for the age squared term was smallest for the DNAmClock_Cortical_ (beta = −0.002, P = 1.94E-07), again highlighting that bespoke clocks can be used to minimise bias in subsequent analyses.

**Table 3 –.** The relationship between DNAm age and age (age and age^2^) using different DNAm clock algorithms. DNAm age was estimated using our novel cortical clock,, Horvath’s multi-tissue clock (Horvath, 2013), Zhang’s elastic net blood clock (Zhang et al., 2019) and Levine’s Pheno Age clock (Levine et al., 2018) in the “testing” dataset (n = 350 cortical samples), the “validation” dataset (n =1221 cortical samples) and the blood dataset (n =1175 whole blood samples).

|  |  | Testing dataset |  |  | Validation dataset |  |  | Blood dataset |  |  |
| --- | --- | --- | --- | --- | --- | --- | --- | --- | --- | --- |
|  |  | Beta | SE | P | Beta | SE | P | Beta | SE | P |
| **Cortical Clock** | **DNAm age vs age** | 1.137 | 0.034 | 2.86E-108 | 1.031 | 0.174 | 5.31E-09 | 0.585 | 0.063 | 5.37E-20 |
|  | **DNAm age vs age<sup>2</sup>** | -0.002 | 0.000 | 1.94E-07 | -0.002 | 0.001 | 0.028 | 0.000 | 0.001 | 0.702 |
| **Multi-tissue clock** | **DNAm age vs age** | 1.082 | 0.041 | 3.17E-83 | 0.683 | 0.164 | 3.51E-05 | 0.754 | 0.065 | 6.01E-30 |
|  | **DNAm age vs age<sup>2</sup>** | -0.004 | 0.000 | 2.45E-21 | -0.002 | 0.001 | 0.085 | -0.001 | 0.001 | 3.71E-02 |
| **Blood Clock** | **DNAm age vs age** | 0.821 | 0.034 | 1.30E-74 | 0.640 | 0.175 | 3.00E-04 | 1.145 | 0.046 | 9.50E-111 |
|  | **DNAm age vs age<sup>2</sup>** | -0.003 | 0.000 | 1.81E-21 | -0.002 | 0.001 | 0.057 | -0.002 | 0.000 | 8.47E-09 |
| **Pheno Age Clock** | **DNAm age vs age** | 0.571 | 0.069 | 3.19E-15 | -0.351 | 0.229 | 0.127 | 0.631 | 0.085 | 1.86E-13 |
|  | **DNAm age vs age<sup>2</sup>** | -0.002 | 0.001 | 4.47E-03 | 0.004 | 0.001 | 0.014 | 0.001 | 0.001 | 0.388 |

### The cortical clock loses accuracy when applied to non-cortical tissues

To assess the specificity of the novel cortex clock we next applied each of the DNAm age clocks to a large whole blood DNAm dataset (n = 1175; age range = 28 - 98 years; mean age = 57.96 years). Although the DNAmClock_Cortical_ actually performed remarkably well on whole blood (r = 0.88), with a similar predictive ability to the DNAmClock_Multi_ (r = 0.90) (**Fig. 3** and **Supplementary Fig. S5**), there was a non-linear relationship between DNAm age and age estimated using this clock and a systematic under prediction of DNAm age in samples from people aged over 60 years (**Fig. 3A**(**i**) and **Fig. 3B**(**i**)). The DNAmClock_Blood_ performed best on the blood dataset (r = 0.97), outperforming the three other clocks (**Table 4; Fig.3; Supplementary Fig. S5 and Supplementary Fig. S6**) and providing further support for the notion that epigenetic clocks work optimally for the tissue-type on which they are calibrated.

**Figure 3:**
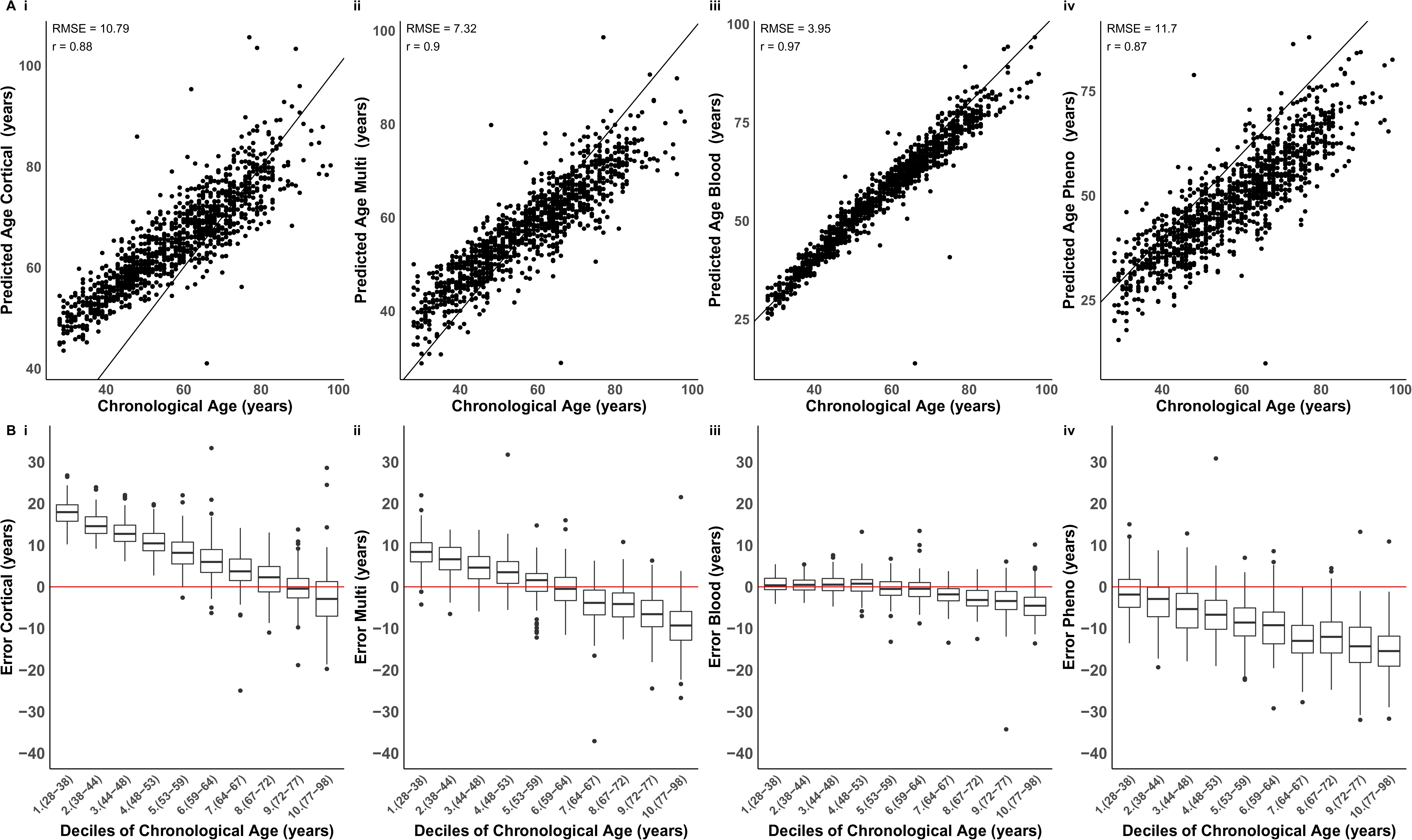
The blood based DNA methylation clock performs best in data derived from whole blood samples. (**A**) Shown is a comparison of DNA methylation age estimates against chronological age in a large whole blood dataset (n = 1175), where DNAm age derived using four DNA methylation age clocks: (**i**) our novel DNAmClock_Cortical_; (**ii**) Horvath’s DNAmClock_Multi_; (**iii**) Zhang’s DNAmClock_Blood_ and (**iv**) Levine’s DNAmClock_Pheno_. The x-axis represents chronological age (years), the y-axis represents predicted age (years). Each point on the plot represents an individual in the whole blood dataset. Our novel clock does not predict as well in the cortex, although it has a similar predictive ability to Horvath’s clock. The distribution of the error (DNA methylation age - chronological age) is presented in (**B**) for each decile for each of the four DNA methylation clocks. Deciles were calculated by assigning chronological age into ten bins and are represented along the x-axis by the numbers one to ten, followed by brackets which display the age range included in each decile. The ends of the boxes are the upper and lower quartiles of the errors, the horizontal line inside the box represents the median deviation and the two lines outside the boxes extend to the highest and lowest observations. Outliers are represented by points beyond these lines. The red horizontal line represents perfect prediction (zero error). *DNAmClock_Cortical_= Cortical DNA methylation age Clock; DNAmClock_Multi_ = Multi-tissue DNA methylation age clock; DNAmClock_Blood_ = Blood DNA methylation age clock and DNAmClock_Pheno_ = Pheno Age DNA methylation age clock.

**Table 4 –.** The cortex clock is less accurate at estimating DNA methylation age algorithms in blood compared to cortex tissue samples. Although still compares well to existing clock algorithms. Accuracy statistics between DNAm age estimates and chronological age using our novel cortical clock, Horvath’s multi-tissue clock (Horvath, 2013), Zhang’s elastic net blood clock (Zhang et al., 2019) and Levine’s Pheno Age clock (Levine et al., 2018) in our blood dataset (n = 1175 whole blood samples). RMSE = root mean squared error (years). MAD = mean absolute deviation (years).

|  | **Cortical Clock** | **Multi-tissue Clock** | **Blood Clock** | **Pheno Age Clock** |
| --- | --- | --- | --- | --- |
| **Correlation (r)** | 0.88 | 0.90 | 0.97 | 0.87 |
| **RMSE (years)** | 10.79 | 7.32 | 3.95 | 11.70 |

## Discussion

Existing DNAm age clocks have been widely utilised for predicting age and exploring accelerated ageing in disease, although there is evidence of systematic underestimation of DNAm age in older samples, particularly in the brain (El Khoury *et al*., 2019). We developed a novel epigenetic age model specifically for human cortex - the cortical DNAm clock (DNAmClock_Cortical_) - built using an extensive collection of DNAm data from >1000 human cortex samples. Our model dramatically outperforms existing DNAm-based biomarkers for age prediction in data derived from the human cortex.

There are several potential causes of the systematic underestimation of DNAm age in the cortex, especially in samples from elderly donors, when using existing DNAm clocks such as Hovath’s DNAmClock_Multi_ (Horvath, 2013), Zhang’s DNAmClock_Blood_ (Zhang *et al*., 2019) and Levine’s DNAmClock_Pheno_ (Levine *et al*., 2018). First, it may be a consequence of the distribution of ages in the training data used in existing clocks; these clocks were derived using samples containing a relatively small proportion of samples from human brain and/or from older people. Second, as there is evidence for cell-type and tissue specific patterns of DNAm (Mendizabal *et al*., 2019), the observed imprecision may reflect a consequence of underfitting the model across tissues. Third, the relationship between DNA methylation and age may not be linear across the lifespan, and a non-linear model is needed to capture attenuated effects in older samples. This would be comparable to the transformation required to accurately predict DNAm age for younger samples (0-20 years), where the association between age and with DNA methylation is of larger magnitude.

Our data suggest that both tissue-specificity and the age of samples included in the training dataset influence the precision of DNAm age estimators, as shown by the increase in accuracy when using our cortical clock relative to existing clocks in human cortex tissue samples. This notion is further supported by the accuracy we found using the blood-based estimators on a large blood dataset. Our observations suggest that tissue type has a major influence on the accuracy of DNAm age clocks, and to accurately predict age it is important to use a clock calibrated specifically for the tissue from which samples have been derived. Our data demonstrate that the performance of existing DNAm clocks varies considerably across ages and is diminished in samples from elderly donors. This is particularly important to consider when assessing DNAm age in the context of diseases and phenotypes that are associated with older age such as dementia and neurodegenerative disease. Our results show that it is important to use a clock that has been trained using samples from the relevant age group; the training data used in the development of the DNAmClock_Cortical_ included a good representation of older samples, meaning it overcomes the systematic underestimation of DNAm age in the elderly that was observed with existing clocks. The importance of developing tissuespecific estimators is supported by other recently developed tissue-specific clocks including DNAm age predictors for whole blood (Zhang *et al*., 2019), human skeletal muscle (Voisin *et al*., 2019) and human bone (Gopalan *et al*., 2019), which all out perform pan-tissue clocks in samples from the specific tissues in which they were trained. It is known that DNA methylation patterns are distinct between tissue and cell types (Mendizabal *et al*., 2019), and it is therefore not surprising that DNAm age estimation models would differ in accuracy across tissue types. As technologies for profiling DNAm in purified cell populations, future clocks should be developed for specific cell-types to overcome issues of cellular heterogeneity in complex tissues such as the brain.

Although a pan-tissue estimator such as Horvath’s DNAmClock_Multi_ has clear general utility, the trade-off between accuracy and practicality needs to be taken into consideration depending on the hypothesised question being tested. Applying one model across multiple tissues may lead to a suboptimal fit (for example, when applying a linear model where there is non-linearity). To assess the linearity of DNAm age predictors we investigated the association between DNAm age, and age squared. Adding the squared variable allowed us to more accurately model the effect of age, which could have a non-linear relationship with DNAm age. The DNAmClock_Cortical_ was the most linear in terms of fitting DNAm age against actual age. Although age squared terms were significantly associated with DNAm age in the testing data using all estimators, the higher significance of the age squared term in the cortex-specific clock suggests that of all the clocks, our model is the least biased. However, as indicated by the relationship between DNAm and age squared, we need to consider the possibility that fitting a linear model might not be the best approach, and to account for this possibility we recommend that future age-acceleration analyses control for age squared terms.

Due to the nature of DNAm clocks, Δ age estimated using existing clocks is highly correlated with chronological age (El Khoury *et al*., 2019). If age is not controlled for it could lead to spurious associations with health outcomes, which are driven by age and not the variable of interest. Furthermore, as the prediction is less precise in older individuals, even where DNAm is regressed on chronological age, the residual may still be associated with age, potentially leading to false positive associations. Recent studies have found associations between accelerated DNAm age in human brain and neurodegenerative phenotypes (Levine *et al*., 2015, 2018). Our findings suggest that previous associations with age-associated phenotypes may have been confounded by a lack of robust calibration to estimate DNAm age in human cortex from elderly donors; caution is warranted in interpreting reported results. While, DNAm age is a useful indicator of age, it may not be the best indicator of health disparities between individuals with brain disorders.

In summary, we show that previous epigenetic clocks systematically underestimate age in older samples and do not perform as well in human cortex tissue. We developed a novel epigenetic age model specifically for human cortex. Our findings suggest that previous associations between predicted DNAm age and neurodegenerative phenotypes may represent false positives resulting from suboptimal calibration of DNAm clocks for the tissue being tested and for phenotypes that manifest at older ages. The age distribution and tissue type of samples included in training datasets need to be considered when building and applying epigenetic clock algorithms to human epidemiological or disease cohorts.

## Acknowledgements

We would like to gratefully acknowledge all donors and their families for the tissue provided for this study. Human post-mortem tissue was obtained from the South West Dementia Brain Bank, London Neurodegenerative Diseases Brain Bank, Manchester Brain Bank, Newcastle Brain Tissue Resource and Oxford Brain Bank, members of the Brains for Dementia Research (BDR) Network. We wish to acknowledge the neuropathologists at each centre and BDR Brain Bank staff for the collection and classification of the samples.

## References

Baker DJ, Wijshake T, Tchkonia T, LeBrasseur NK, Childs BG, van de Sluis B, et al. Clearance of p16Ink4a-positive senescent cells delays ageing-associated disorders. Nature 2011; 479: 232–236.

Bell JE, Alafuzoff I, Al-Sarraj S, Arzberger T, Bogdanovic N, Budka H, et al. Management of a twenty-first century brain bank: experience in the BrainNet Europe consortium. Acta Neuropathol. 2008; 115: 497–507.

Bernstein BE, Meissner A, Lander ES. The mammalian epigenome. Cell 2007; 128: 669–681.

Buck N, McFall S. Understanding Society: design overview. Longitudinal and Life Course Studies 2011

Campisi J, Vijg J. Does damage to DNA and other macromolecules play a role in aging? If so, how? J. Gerontol. A, Biol. Sci. Med. Sci. 2009; 64: 175–178.

Chen Y, Lemire M, Choufani S, Butcher DT, Grafodatskaya D, Zanke BW, et al. Discovery of cross-reactive probes and polymorphic CpGs in the Illumina Infinium HumanMethylation450 microarray. Epigenetics 2013; 8: 203–209.

Chouliaras L, Pishva E, Haapakoski R, Zsoldos E, Mahmood A, Filippini N, et al. Peripheral DNA methylation, cognitive decline and brain aging: pilot findings from the Whitehall II imaging study. Epigenomics 2018; 10: 585–595.

Chuang Y-H, Paul KC, Bronstein JM, Bordelon Y, Horvath S, Ritz B. Parkinson’s disease is associated with DNA methylation levels in human blood and saliva. Genome Med. 2017; 9: 76.

El Khoury LY, Gorrie-Stone T, Smart M, Hughes A, Bao Y, Andrayas A, et al. Systematic underestimation of the epigenetic clock and age acceleration in older subjects. Genome Biol. 2019; 20: 283.

Elliott HR, Tillin T, McArdle WL, Ho K, Duggirala A, Frayling TM, et al. Differences in smoking associated DNA methylation patterns in South Asians and Europeans. Clin. Epigenetics 2014; 6: 4.

Francis PT, Costello H, Hayes GM. Brains for dementia research: evolution in a longitudinal brain donation cohort to maximize current and future value. J. Alzheimers Dis. 2018; 66: 1635–1644.

Friedman J, Hastie T, Tibshirani R. Regularization Paths for Generalized Linear Models via Coordinate Descent. J Stat Softw 2010; 33: 1–22.

Gopalan S, Gaige J, Henn BM. DNA methylation-based forensic age estimation in human bone. BioRxiv 2019

Gorrie-Stone TJ, Smart MC, Saffari A, Malki K, Hannon E, Burrage J, et al. Bigmelon: tools for analysing large DNA methylation datasets. Bioinformatics 2019; 35: 981–986.

Hannon E, Gorrie-Stone TJ, Smart MC, Burrage J, Hughes A, Bao Y, et al. Leveraging DNA-Methylation Quantitative-Trait Loci to Characterize the Relationship between Methylomic Variation, Gene Expression, and Complex Traits. Am. J. Hum. Genet. 2018; 103: 654–665.

Hannum G, Guinney J, Zhao L, Zhang L, Hughes G, Sadda S, et al. Genome-wide methylation profiles reveal quantitative views of human aging rates. Mol. Cell 2013; 49: 359–367.

Harper S. Economic and social implications of aging societies. Science 2014; 346: 587–591.

Horvath S, Oshima J, Martin GM, Lu AT, Quach A, Cohen H, et al. Epigenetic clock for skin and blood cells applied to Hutchinson Gilford Progeria Syndrome and ex vivo studies. Aging (Albany, NY) 2018; 10: 1758–1775.

Horvath S, Ritz BR. Increased epigenetic age and granulocyte counts in the blood of Parkinson’s disease patients. Aging (Albany, NY) 2015; 7: 1130–1142.

Horvath S, Zhang Y, Langfelder P, Kahn RS, Boks MPM, van Eijk K, et al. Aging effects on DNA methylation modules in human brain and blood tissue. Genome Biol. 2012; 13: R97.

Horvath S. DNA methylation age of human tissues and cell types. Genome Biol. 2013; 14: R115.

Jaffe AE, Gao Y, Deep-Soboslay A, Tao R, Hyde TM, Weinberger DR, et al. Mapping DNA methylation across development, genotype and schizophrenia in the human frontal cortex. Nat. Neurosci. 2016; 19: 40–47.

De Jager PL, Srivastava G, Lunnon K, Burgess J, Schalkwyk LC, Yu L, et al. Alzheimer’s disease: early alterations in brain DNA methylation at ANK1, BIN1, RHBDF2 and other loci. Nat. Neurosci. 2014; 17: 1156–1163.

Jylhävä J, Jiang M, Foebel AD, Pedersen NL, Hägg S. Can markers of biological age predict dependency in old age? Biogerontology 2019; 20: 321–329.

Jylhävä J, Pedersen NL, Hägg S. Biological Age Predictors. EBioMedicine 2017; 21: 29–36.

Knight AK, Craig JM, Theda C, Bækvad-Hansen M, Bybjerg-Grauholm J, Hansen CS, et al. An epigenetic clock for gestational age at birth based on blood methylation data. Genome Biol. 2016; 17: 206.

Levine ME, Lu AT, Bennett DA, Horvath S. Epigenetic age of the pre-frontal cortex is associated with neuritic plaques, amyloid load, and Alzheimer’s disease related cognitive functioning. Aging (Albany, NY) 2015; 7: 1198–1211.

Levine ME, Lu AT, Quach A, Chen BH, Assimes TL, Bandinelli S, et al. An epigenetic biomarker of aging for lifespan and healthspan. Aging (Albany, NY) 2018; 10: 573–591.

Lunnon K, Smith R, Hannon E, De Jager PL, Srivastava G, Volta M, et al. Methylomic profiling implicates cortical deregulation of ANK1 in Alzheimer’s disease. Nat. Neurosci. 2014; 17: 1164–1170.

Marioni RE, Shah S, McRae AF, Ritchie SJ, Muniz-Terrera G, Harris SE, et al. The epigenetic clock is correlated with physical and cognitive fitness in the Lothian Birth Cohort 1936. Int. J. Epidemiol. 2015; 44: 1388–1396.

McCartney DL, Stevenson AJ, Walker RM, Gibson J, Morris SW, Campbell A, et al. Investigating the relationship between DNA methylation age acceleration and risk factors for Alzheimer’s disease. Alzheimers Dement (Amst) 2018; 10: 429–437.

McEwen LM, O’Donnell KJ, McGill MG, Edgar RD, Jones MJ, MacIsaac JL, et al. The PedBE clock accurately estimates DNA methylation age in pediatric buccal cells. Proc. Natl. Acad. Sci. USA 2019

McKinney BC, Lin H, Ding Y, Lewis DA, Sweet RA. DNA methylation age is not accelerated in brain or blood of subjects with schizophrenia. Schizophr. Res. 2018; 196: 39–44.

Mendizabal I, Berto S, Usui N, Toriumi K, Chatterjee P, Douglas C, et al. Cell type-specific epigenetic links to schizophrenia risk in the brain. Genome Biol. 2019; 20: 135.

Mendizabal I, Yi SV. Whole-genome bisulfite sequencing maps from multiple human tissues reveal novel CpG islands associated with tissue-specific regulation. Hum. Mol. Genet. 2016; 25: 69–82.

Moran S, Arribas C, Esteller M. Validation of a DNA methylation microarray for 850,000 CpG sites of the human genome enriched in enhancer sequences. Epigenomics 2016; 8: 389–399.

Oberdoerffer P, Sinclair DA. The role of nuclear architecture in genomic instability and ageing. Nat. Rev. Mol. Cell Biol. 2007; 8: 692–702.

Pidsley R, Viana J, Hannon E, Spiers H, Troakes C, Al-Saraj S, et al. Methylomic profiling of human brain tissue’’ supports a neurodevelopmental origin for schizophrenia. Genome Biol. 2014; 15: 483.

Pidsley R, Y Wong CC, Volta M, Lunnon K, Mill J, Schalkwyk LC. A data-driven approach to preprocessing Illumina 450K methylation array data. BMC Genomics 2013; 14: 293.

Samarasekera N, Al-Shahi Salman R, Huitinga I, Klioueva N, McLean CA, Kretzschmar H, et al. Brain banking for neurological disorders. Lancet Neurol. 2013; 12: 1096–1105.

Sanders JL, Newman AB. Telomere length in epidemiology: a biomarker of aging, age-related disease, both, or neither? Epidemiol Rev 2013; 35: 112–131.

Sierra F. Geroscience and the challenges of aging societies. Aging Med. 2019; 2: 132–134.

Simpkin AJ, Suderman M, Howe LD. Epigenetic clocks for gestational age: statistical and study design considerations. Clin. Epigenetics 2017; 9: 100.

Smith AR, Smith RG, Condliffe D, Hannon E, Schalkwyk L, Mill J, et al. Increased DNA methylation near TREM2 is consistently seen in the superior temporal gyrus in Alzheimer’s disease brain. Neurobiol. Aging 2016; 47: 35–40.

Smith AR, Smith RG, Pishva E, Hannon E, Roubroeks JAY, Burrage J, et al. Parallel profiling of DNA methylation and hydroxymethylation highlights neuropathology-associated epigenetic variation in Alzheimer’s disease. Clin. Epigenetics 2019; 11: 52.

Smith RG, Hannon E, De Jager PL, Chibnik L, Lott SJ, Condliffe D, et al. Elevated DNA methylation across a 48-kb region spanning the HOXA gene cluster is associated with Alzheimer’s disease neuropathology. Alzheimers Dement. 2018; 14: 1580–1588.

Sosnoff JJ, Newell KM. Are age-related increases in force variability due to decrements in strength? Exp. Brain Res. 2006; 174: 86–94.

Sugden K, Hannon EJ, Arseneault L, Belsky DW, Broadbent JM, Corcoran DL, et al. Establishing a generalized polyepigenetic biomarker for tobacco smoking. Transl. Psychiatry 2019; 9: 92.

Voisin S, Harvey NR, Haupt LM, Griffiths LR, Ashton KJ, Coffey VG, et al. An epigenetic clock for skeletal muscle. BioRxiv 2019

Wong CCY, Smith RG, Hannon E, Ramaswami G, Parikshak NN, Assary E, et al. Genome-wide DNA methylation profiling identifies convergent molecular signatures associated with idiopathic and syndromic autism in post-mortem human brain tissue. Hum. Mol. Genet. 2019; 28: 2201–2211.

Yu L, Chibnik LB, Srivastava GP, Pochet N, Yang J, Xu J, et al. Association of Brain DNA methylation in SORL1, ABCA7, HLA-DRB5, SLC24A4, and BIN1 with pathological diagnosis of Alzheimer disease. JAMA Neurol. 2015; 72: 15–24.

Zhang Q, Vallerga CL, Walker RM, Lin T, Henders AK, Montgomery GW, et al. Improved precision of epigenetic clock estimates across tissues and its implication for biological ageing. Genome Med. 2019; 11: 54.

Zou H, Hastie T. Regularization and variable selection via the elastic net. J Royal Statistical Soc B 2005

