## Supplementary Figures for "Recalibrating the Epigenetic Clock: Implications for Assessing Biological Age in the Human Cortex"

* These authors contributed equally.

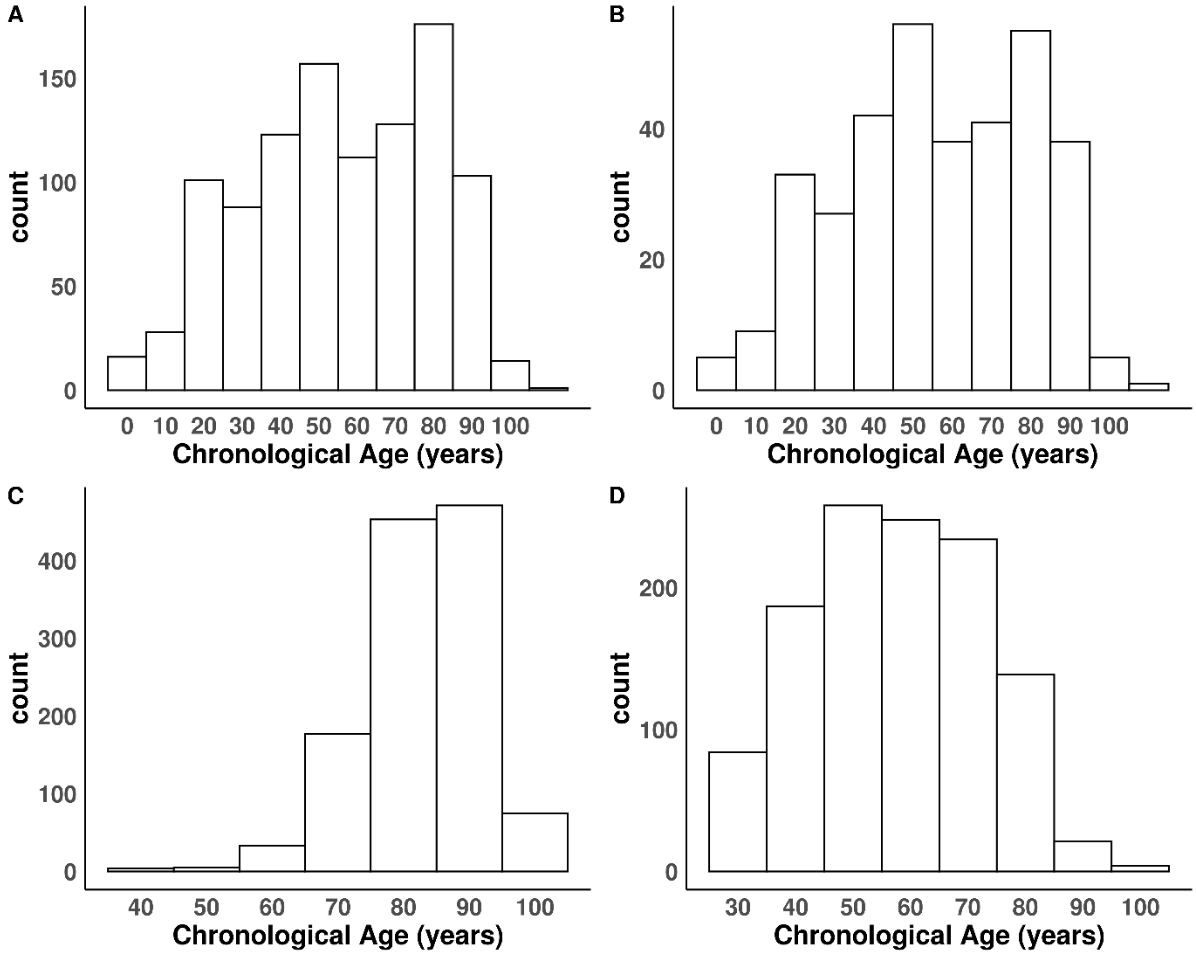


**Figure S1: Histograms showing the distribution of chronological age in the datasets used in the study.** (***A***) *The training dataset (n = 1,047 cortical samples);* (***B***) *the testing dataset (n = 350 cortical samples);* (***C***) *the validation dataset (n = 1221 cortical samples) and* (***D***) *the whole blood dataset (n = 1175 whole blood samples).*

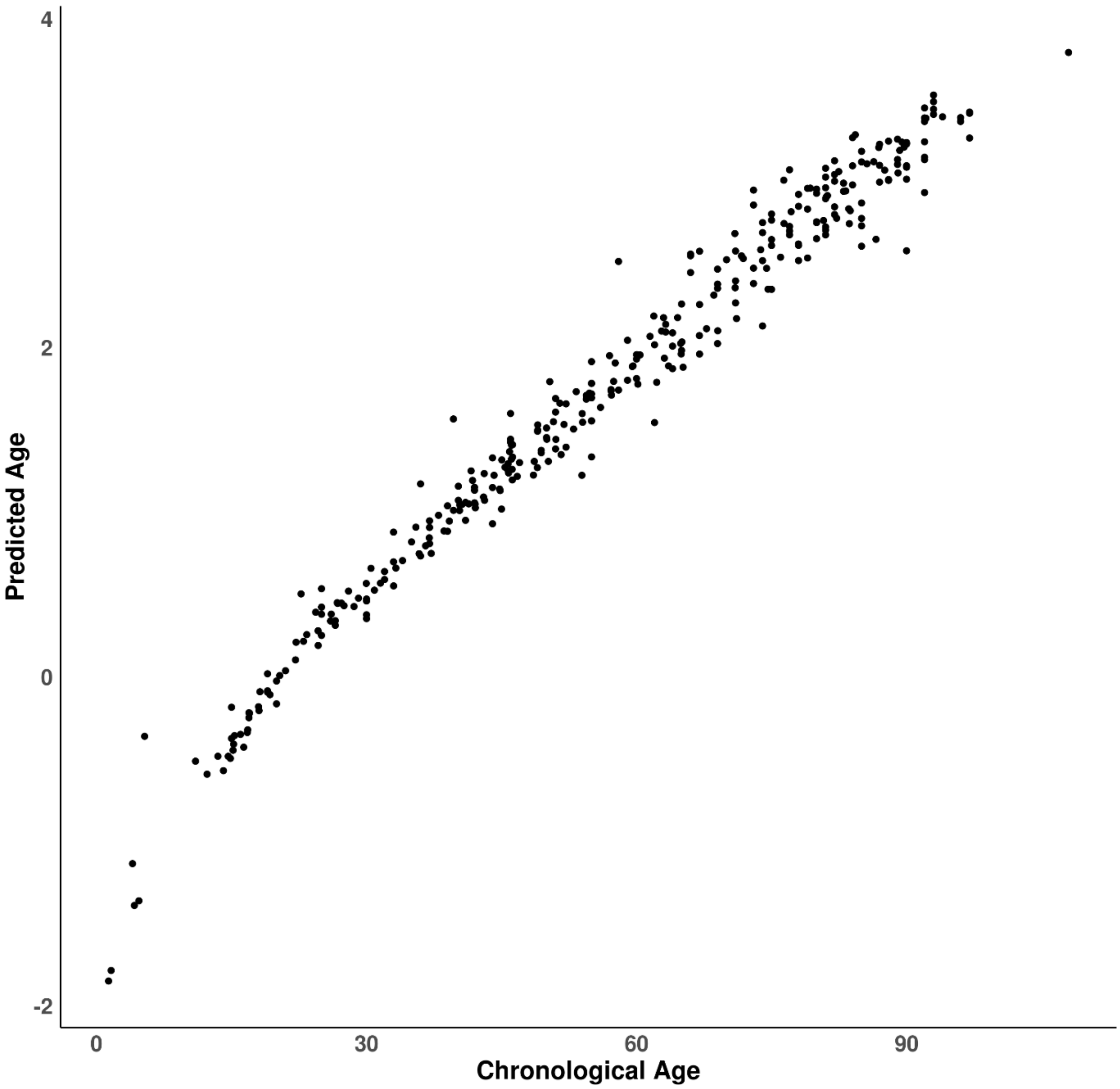


**Figure S2: DNA methylation age has a logarithmic relationship with chronological age between the ages of 0-20 years**. From 20 years onward there is a linear relationship with chronological age. The x-axis represents chronological age (years), the y-axis represents predicted age prior to applying the anti-transformation function, whereby age between 0-20 years is log transformed, and ages 20+ are transformed to account for this. Each point on the plot is a sample from the testing dataset (n = 350 cortical samples).

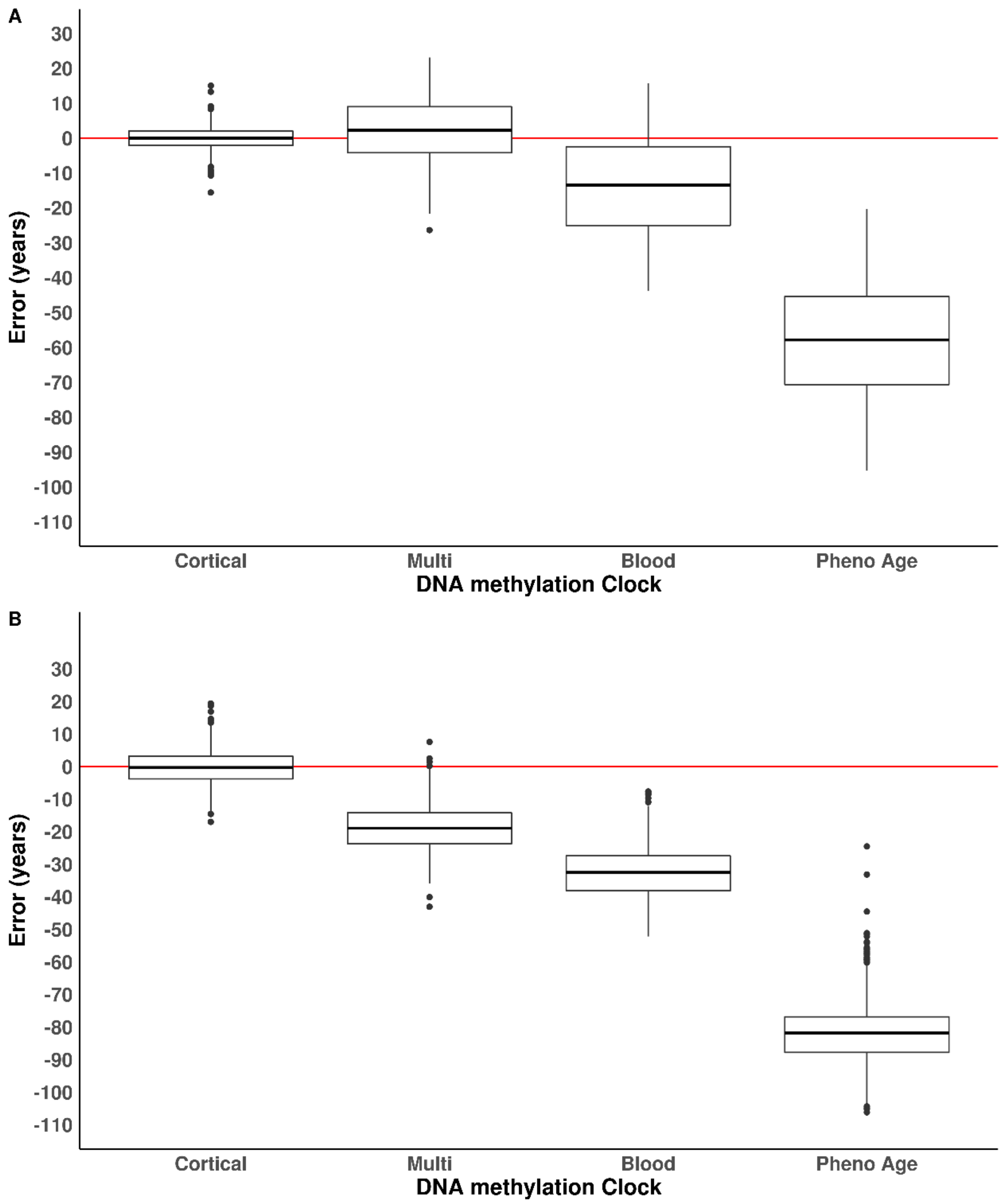


**Figure S3: The cortical DNAm age clock has elevated accuracy in human cortex samples compared to existing DNAm clocks.** Shown is the distribution of the error (DNA methylation age - chronological age) for each of the four DNA methylation age clocks in (**A**) the testing dataset (n = 350 cortical samples) and (**B**) the validation dataset (n = 1221 cortical samples). The ends of the boxes are the upper and lower quartiles of the errors, the horizontal line inside the box represents the median deviation and the two lines outside the boxes extend to the highest and lowest observations. Outliers are represented by points beyond these lines. The red horizontal line represents perfect prediction (zero error).

*$DNAmClock_{Cortical}$ = Cortical DNA methylation age Clock; $DNAmClock_{Multi}$ = Multi-tissue DNA methylation age clock;$DNAmClock_{Blood}$ = Blood DNA methylation age clock and $DNAmClock_{Pheno}$ = Pheno Age DNA methylation age clock.

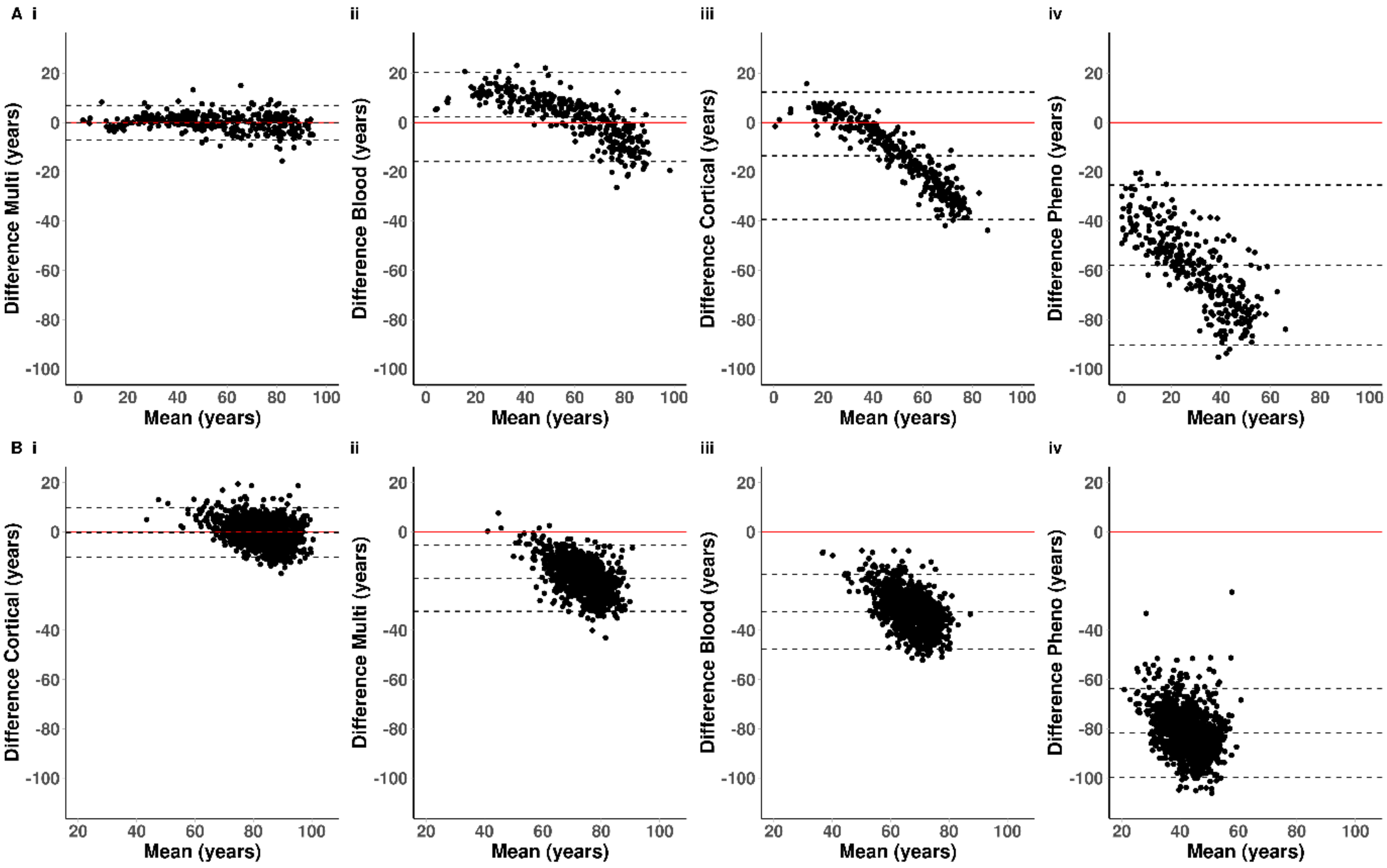

A i
ii
iii
iv
Difference Multi (years)
Difference Blood (years)
Difference Cortical (years)
Difference Pheno (years)
Mean (years)
B i
Difference Cortical (years)
Difference Multi (years)
Difference Blood (years)
Difference Pheno (years)

**Figure S4: Bland-Altman plots highlighting elevated performance of the cortical DNAm clock in human cortex tissue across the lifespan**. Shown is the mean difference between actual chronological age and estimated DNAm ages derived in (**A**) the testing dataset (n= 350 cortical samples) and (**B**) the validation dataset (n = 1221 cortical samples), where DNAm age derived using four DNA methylation age clocks: (**i**) our novel $DNAmClock_{Cortical}$; (**ii**) Horvath's $DNAmClock_{Multi}$; (**iii**) Zhang's $DNAmClock_{Blood}$ and (**iv**) Levine's $DNAmClock_{Pheno}$ The dashed horizontal lines in each case are the mean difference and +/- 1.96 * standard deviation; for normally distributed difference due to error 5% points would lie outside these. The solid horizontal line represents where the mean difference would be zero.

*$DNAmClock_{Cortical}$ = Cortical DNA methylation age Clock; $DNAmClock_{Multi}$ = Multi-tissue DNA methylation age clock; $DNAmClock_{Blood}$ = Blood DNA methylation age clock and $DNAmClock_{Pheno}$ = Pheno Age DNA methylation age clock.

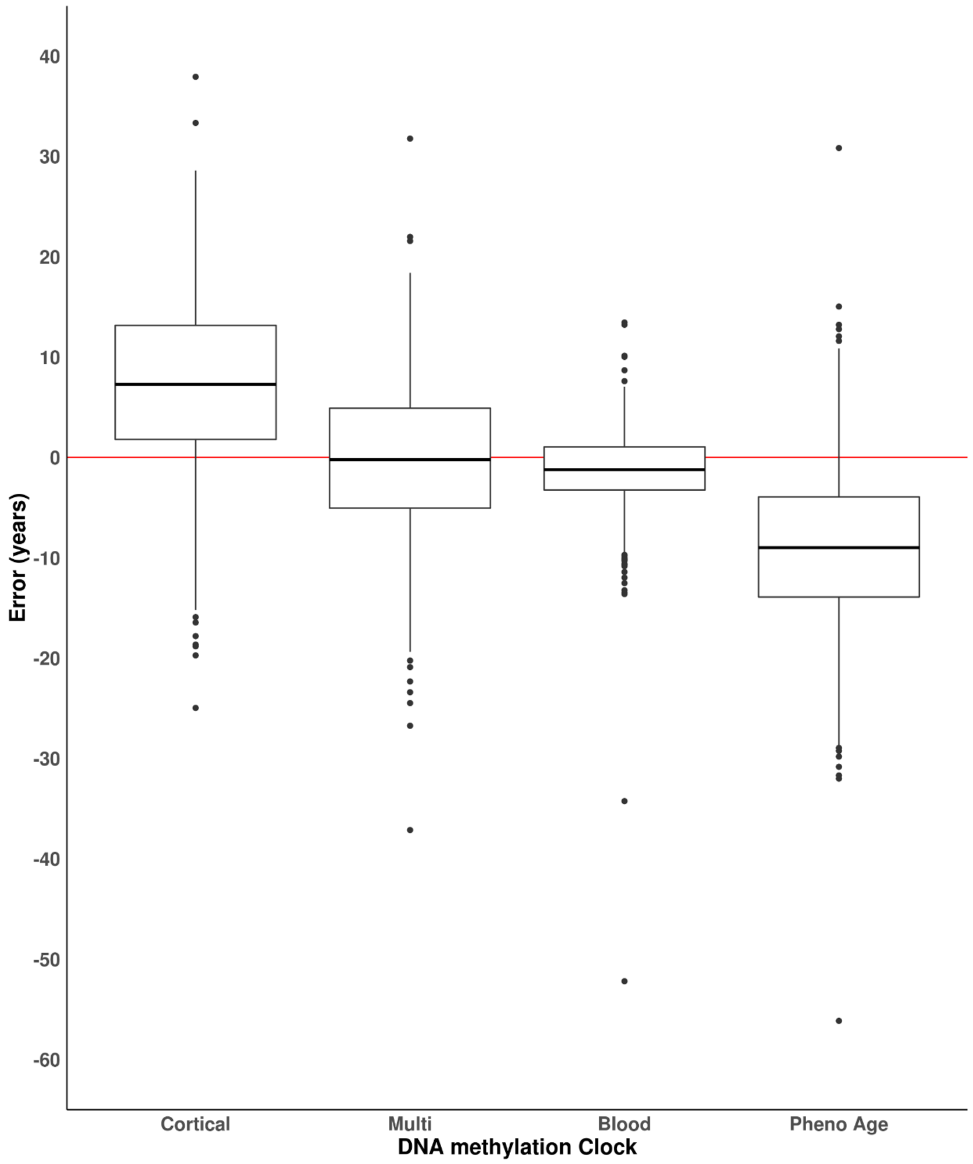


**Figure S5**: ***The blood DNAm age clock has elevated accuracy in human whole blood samples* compared to non-tissue specific DNAm clocks.** Distribution of the error in years (DNA methylation age - chronological age) comparing four DNA methylation age clocks: our novel $DNAmClock_{Cortical}$, the $DNAmClock_{Multi}$, the $DNAmClock_{Blood}$ and the $DNAmClock_{Pheno}$ in the **whole blood dataset** (n = 1175). The ends of the boxes are the upper and lower quartiles of the errors, the horizontal line inside the box represents the mean absolute deviation and the two lines outside the boxes extend to the highest and lowest observations. Outliers are represented by points beyond these lines.

*$DNAmClock_{Cortical}$ = Cortical DNA methylation age Clock; $DNAmClock_{Multi}$ = Multi-tissue DNA methylation age clock; $DNAmClock_{Blood}$ = Blood DNA methylation age clock and $DNAmClock_{Pheno}$ = Pheno Age DNA methylation age clock.

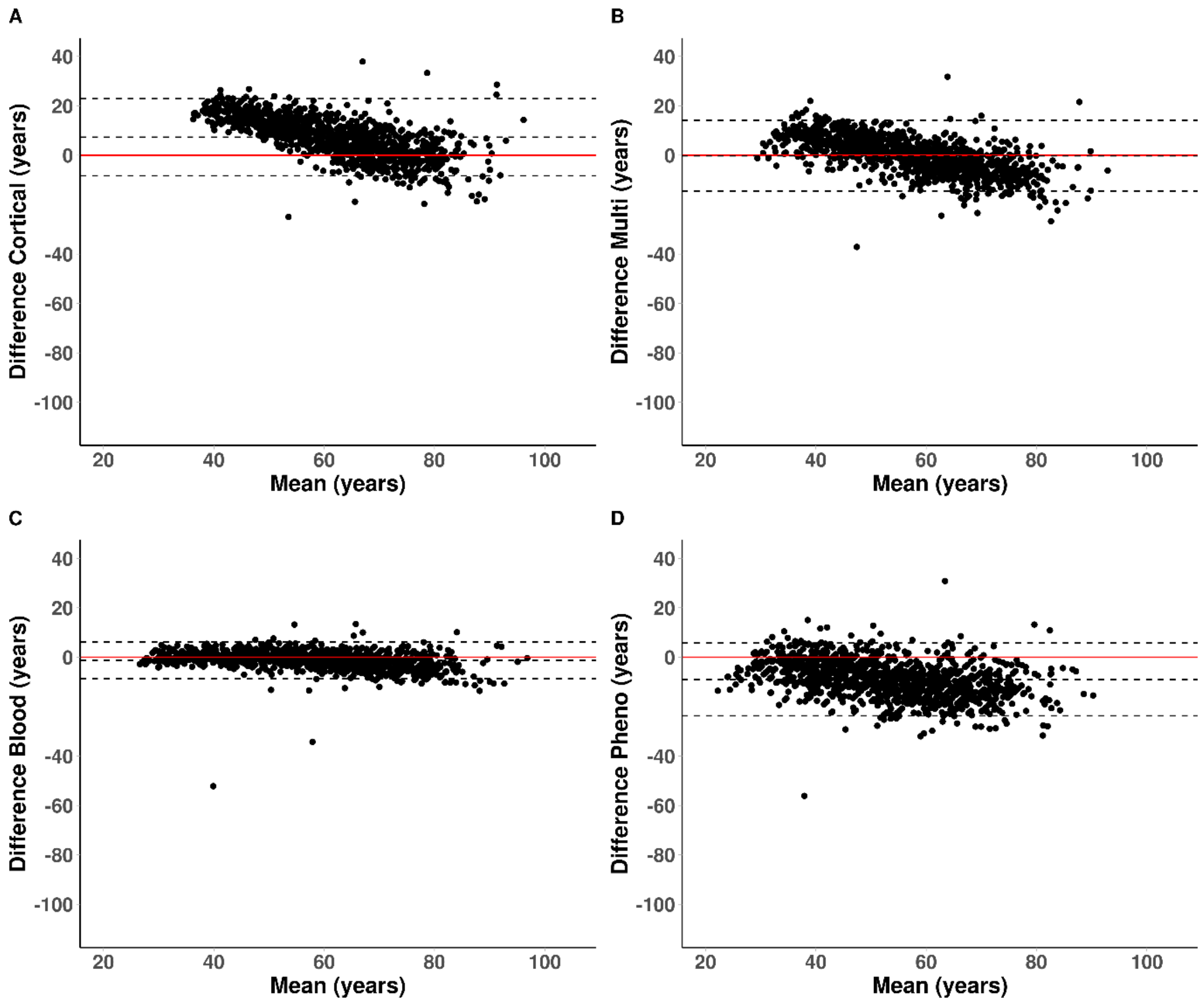


**Figure S6**: **Bland-Altman plots highlighting elevated performance of blood based DNAm clocks in whole blood samples.** *Mean-difference (Bland-Altman) plots showing the difference between DNA methylation age estimates against chronological age using* (**A**) *our novel* $\text{DNAmClock}_{\text{Cortical}}$, (**B**) the $\text{DNAmClock}_{\text{Multi}}$, (**C**) the $\text{DNAmClock}_{\text{Blood}}$ and (**D**) the $\text{DNAmClock}_{\text{Pheno}}$ *in the whole blood cohort (n = 1175). The dashed horizontal lines in each case are the mean difference and +/- 1.96 * standard deviation; for normally distributed difference due to error 5% of points would lie outside these. The solid horizontal line represents where the mean difference would be zero.*

*$\text{DNAmClock}_{\text{Cortical}}$ = Cortical DNA methylation age Clock; $\text{DNAmClock}_{\text{Multi}}$ = Multi-tissue DNA methylation age clock; $\text{DNAmClock}_{\text{Blood}}$ = Blood DNA methylation age clock and $\text{DNAmClock}_{\text{Pheno}}$ = Pheno Age DNA methylation age clock.
